# IGF2BP1/IMP1 deletion enhances a facultative stem cell state via regulation of *MAP1LC3B*

**DOI:** 10.1101/2022.02.28.482365

**Authors:** Louis R. Parham, Patrick A. Williams, Priya Chatterji, Kofi K. Acheampong, Charles H. Danan, Kay Katada, Xianghui Ma, Lauren A. Simon, Kaitlyn E. Naughton, Rei Mizuno, Tatiana Karakasheva, Emily A. McMillan, Kelly A. Whelan, Donita C. Brady, Sydney M. Shaffer, Kathryn E. Hamilton

**Affiliations:** Division of Gastroenterology, Hepatology, and Nutrition; Department of Pediatrics; Children’s Hospital of Philadelphia; University of Pennsylvania Perelman School of Medicine, Philadelphia, PA, USA; Kojin Therapeutics, Inc, Boston, MA, USA; Department of Pathology and Laboratory Medicine, University of Pennsylvania Perelman School of Medicine, Philadelphia, PA, USA; Department of Surgery, Uji-Tokushukai Medical Center, Uji, Kyoto, Japan; Department of Pathology & Laboratory Medicine, Lewis Katz School of Medicine at Temple University, Philadelphia, PA, USA; Fels Institute for Cancer Research & Molecular Biology, Lewis Katz School of Medicine at Temple University, Philadelphia, PA, USA; Department of Cancer Biology, Perelman School of Medicine, University of Pennsylvania, Philadelphia, PA, USA; Abramson Family Cancer Research Institute, Perelman School of Medicine, University of Pennsylvania, Philadelphia, PA, USA; Department of Bioengineering, University of Pennsylvania, Philadelphia, PA, USA; Institute for Regenerative Medicine, University of Pennsylvania, Philadelphia, PA, USA

## Abstract

Homeostatic tissue maintenance requires coordinated regulation of metabolic processes including macroautophagy/autophagy. Autophagy dysregulation underlies numerous human diseases. Our prior work revealed that the RNA binding protein IGF2BP1/IMP1 binds transcripts encoding autophagy-related proteins. Furthermore, Imp1 deletion in gastrointestinal epithelial cells in mice was associated with enhanced autophagy flux and improved recovery from tissue injury. In the current study, we evaluated molecular mechanisms underlying IMP1 modulation of autophagy. We provide a mechanism of direct IMP1 regulation of MAP1LC3B that is dependent upon IMP1 phosphorylation or cell stress, suggesting dynamic modulation of Imp1-mediated autophagy repression that facilitates tissue regeneration. More broadly, our study supports a new mechanism by which tissue regeneration is modulated post-transcriptionally via cell state rather than changes in stem or other cell lineages. This new mechanism may be particularly important in gastrointestinal epithelial cells, where autophagy is essential for tissue recovery following injury, or in diseases such as inflammatory bowel disease where defective autophagy is implicated.

---

Lgr5+ active intestinal stem cells (a-ISCs) are the workhorse cells of the epithelium, ensuring a robust pool of cells to execute digestive and barrier functions in the gut [1]. However, a-ISCs are highly susceptible to dying during tissue injury [2]. When a-ISCs are depleted, the epithelium can recover via mobilization of facultative intestinal stem cells (f-ISCs) [2, 3]. F-ISCs are slow-cycling and relatively injury-resistant, and our recent work identified the cell state of high autophagy as a marker of f-ISCs exhibiting DNA damage-resistance and plasticity [4]. In the current study, we sought to elucidate a key regulatory factor of autophagic state in the intestine.

RNA-binding proteins (RBPs) regulate mRNA processing, and ultimately cell state, by governing interactions of mRNAs with the translation machinery, cytoskeletal network, or other trans-acting factors [5]. We demonstrated previously that intestinal epithelial cell-specific deletion of the RNA binding protein Imp1/Igf2bp1 (*Imp1*^*ΔIEC*^) is associated with increased autophagic flux and increased intestinal regeneration following irradiation in *Imp1*^*ΔIEC*^ mice [6, 7]. *Imp1* deletion in *Hopx-CreER*-expressing f-ISCs recapitulated this phenotype, highlighting a specific role for *Imp1* in regulating f-ISC post-irradiation [7]. These data support the hypothesis that Imp1 can influence the census of f-ISCs via modulation of autophagy; however, direct mechanisms remain unknown.

Enhanced regeneration following irradiation suggests increased stem cell capacity in *Imp1*^*ΔIEC*^ mice. Consistent with this observation, we found that cells from *Imp1*^*ΔIEC*^ mice form organoids at approximately double the rate of *Imp1*^*WT*^ mice (Fig. 1A-B). To determine if this phenotype correlated with a shift in f-ISCs, we quantified f-ISCs in *Hopx-CreER;LSL-tdTomato;Imp1*^*fl/fl*^ mice but did not observe a significant difference compared to *Imp1*^*WT*^ cells (Fig. S1A-C). Prior literature suggests that f-ISCs reside within secretory lineages [8-10]. We therefore quantified CD24^high^ (marker of Paneth and enteroendocrine cells) and UEA-1+ (marker of goblet and Paneth cells) cells in *Imp1*^*ΔIEC*^ mice, and similarly did not observe a difference between *Imp1*^*ΔIEC*^ and *Imp1*^*WT*^ mice (Fig. S1D-G). Finally, we evaluated how *Imp1* deletion affects Lgr5-GFP+ a-ISCs in *Imp1*^*ΔIEC*^ mice. Lgr5-GFP;*Imp1*^*ΔIEC*^ mice exhibited a significant reduction of Lgr5-GFP+ cells compared to Lgr5-GFP; *Imp1*^*WT*^ mice (Fig. 1C-E). In summary, we observed an increase in organoid formation in *Imp1*^*ΔIEC*^ mice that is not explained by shifts in f-ISC or a-ISC numbers.

**Figure 1.**
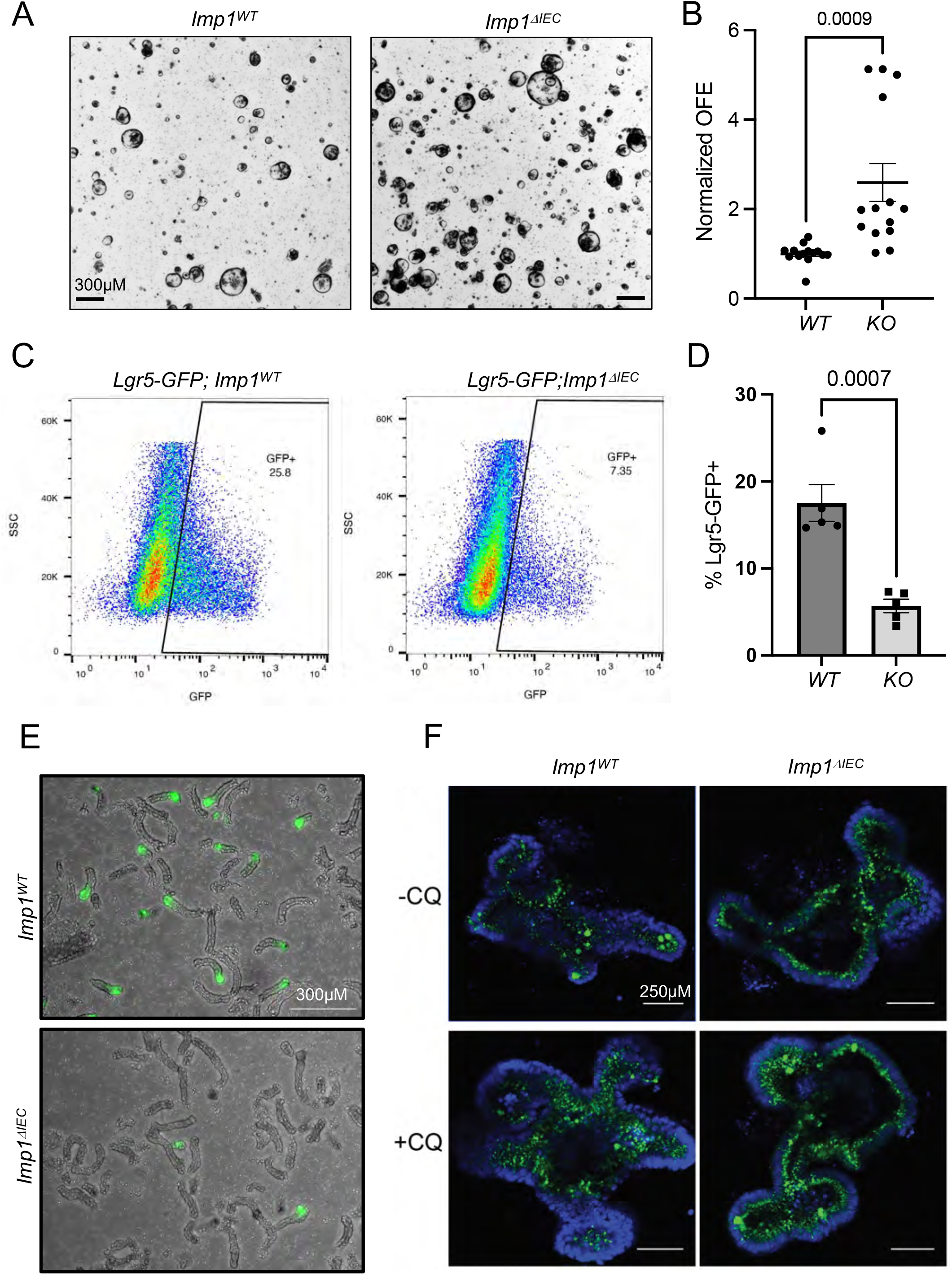
Imp1 deletion enhances organoid formation and autophagic vesicle content. A) Brightfield imaging demonstrating organoid formation efficiency (OFE) in Imp1^WT^ and Imp1^ΔIEC^ mice. B)Quantification of organoid formation efficiency in Imp1^WT^ and Imp1^ΔIEC^ mice. N=3 mice/group, 4-6 wells/mouse. P-value generated by two-tailed student’s t-test. C)Plots demonstrating Lgr5-GFP+ cells in Lgr5-GFP;Imp1^WT^ and Lgr5-GFP;Imp1^ΔIEC^ mice. D)Quantification of mice in (C). N=5 mice/group. P-value generated by two-tailed student’s t-test. E)Representative image of GFP+ crypts from mice in (C). F)Representative CytoID staining in Imp1^WT^ and Imp1^ΔIEC^ enteroids ± chloroquine. N=3 mice/group.

We demonstrated recently that autophagic vesicle levels in IECs correlates with organoid formation ability and injury resistance [4]. We confirmed that *Imp1*^*ΔIEC*^ mice exhibit elevated autophagy in IECs by staining enteroids with the autophagic vesicle marker CytoID. We observed increased CytoID+ puncta in *Imp1*^*ΔIEC*^ compared to *Imp1*^*WT*^ enteroids, which was further enhanced with the autophagy inhibitor chloroquine (Fig. 1F). To ascertain whether enhanced regeneration in *Imp1*^*ΔIEC*^ mice requires autophagy, we compared *Imp1*^*ΔIEC*^ mice to mice with combined deletion of *Imp1* and *Atg7* following irradiation. Autophagy disruption was confirmed via LC3 western blot, which showed the absence of the autophagic vesicle-specific isoform of LC3 known as LC3-II (Fig. S1H). *Imp1*^*ΔIEC*^ mice exhibited an elevated number of regenerative crypt foci, which was abrogated in *Imp1*^*ΔIEC*^*;Atg7*^*ΔIEC*^ mice (Fig. 2A-B). Taken together, these data support the conclusion that enhanced regeneration in *Imp1*^*ΔIEC*^ mice is dependent upon autophagy (Fig. 2A-B).

**Figure 2.**
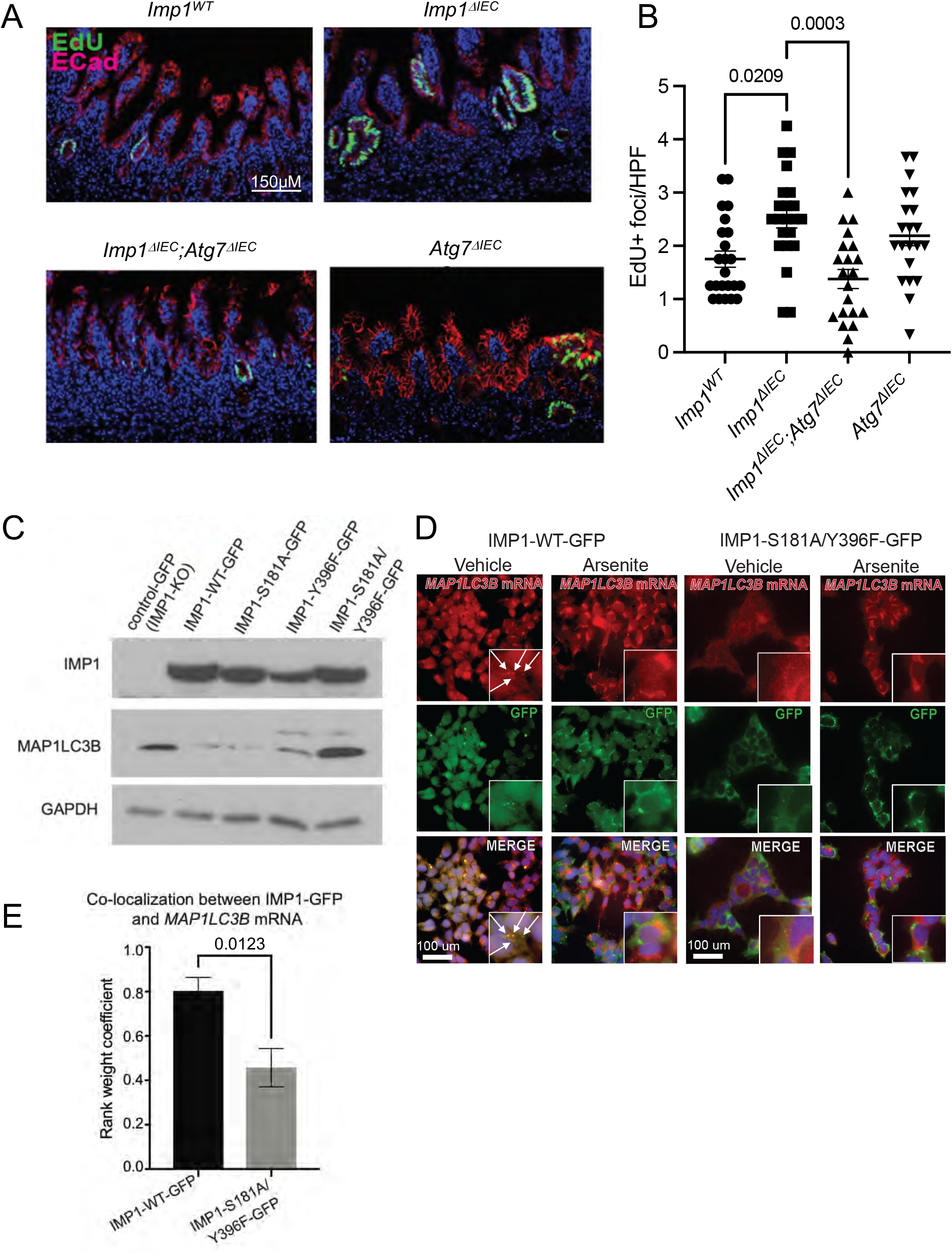
Imp1 modulates intestinal regeneration in an autophagy-dependent manner. A)Representative images of EdU+ foci in indicated mice. B)Quantification of EdU+ foci per high powered field (HPF) in indicated mice. N=4 mice per genotype, ≥20 HPF per animal. P-values generated via one-way ANOVA with Tukey’s multiple comparisons. C)Western blot in IMP1-KO cells with stable expression of indicated GFP constructs. D)Immunofluorescence and smFISH for IMP1-WT-GFP (green) and MAP1LC3B mRNA (red), respectively. White arrows= IMP1-MAP1LC3B co-localization. E)Rank weight coefficient analysis of MAP1LC3 transcripts and IMP1-GFP in indicated cells. P-value generated by two-tailed student’s t-test.

Our prior work showed that IMP1 can bind directly to the autophagy transcript *MAP1LC3B* [6]. We next evaluated the mechanistic contributions of previously described Imp1 phosphorylation sites serine 181 (S181) and tyrosine 396 (Y396), which can affect translation and/or localization of IMP1 targets. We generated *IMP1-KO* HEK293 cell lines with *IMP1-WT-GFP* or phospho-mutants *IMP1-S181A/Y396F-GFP* to evaluate IMP1 co-localization with *MAP1LC3B* transcripts using single molecule mRNA FISH and immunofluorescent imaging of GFP. MAP1LC3B protein expression was increased in control-GFP (IMP1-KO), and IMP1-S181A/Y396F-GFP mutant cells compared to IMP1-WT-GFP (Fig. 2C). IMP1-WT-GFP co-localized with *MAP1LC3B* transcripts (Fig. 2D, white arrows) and co-localization was significantly reduced in IMP1-S181A/Y396F-GFP cells (Fig. 2D) as quantified via rank weight coefficient (Fig 2E). Cell stress induction via arsenite treatment was also associated with reduced co-localization of *MAP1LC3B* mRNA and IMP1-WT-GFP (Fig. 2D). Taken together, our data show that IMP1 co-localizes with *MAP1LC3B* transcripts, supporting a model in which IMP1 can repress autophagy at homeostasis and where de-repression can occur in response to IMP1 phosphorylation and/or cell stress.

In summary, our data support a role for Imp1 regulation of stem cell state via autophagy modulation. We provide a mechanism of direct IMP1 regulation of MAP1LC3B that is dependent upon IMP1 phosphorylation or cell stress, suggesting dynamic modulation of Imp1-mediated autophagy repression that facilitates tissue regeneration. More broadly, our study supports a new mechanism by which tissue regeneration is modulated post-transcriptionally via cell state rather than changes in stem or other cell lineages.

## Supplemental figure legends

**Figure S1.**
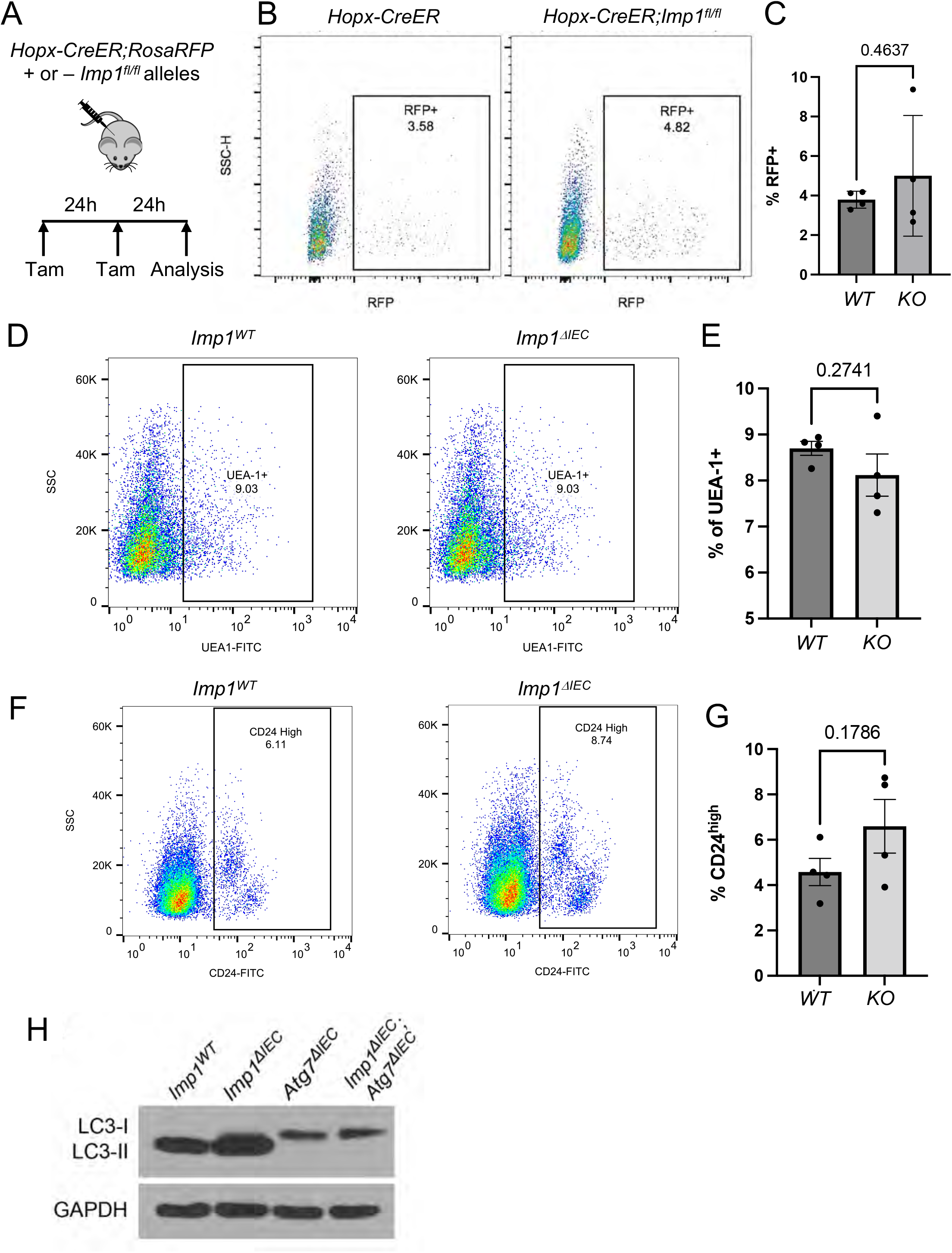
Imp1 deletion does not alter secretory cell or f-ISC frequency at homeostasis. A)Schematic demonstrating tamoxifen scheme for evaluating f-ISC numbers in Hopx-CreER;Rosa-tdTomato mice with and without floxed Imp1 alleles. B)Representative flow plots of RFP+ cells in Hopx-CreER;Rosa-tdTomato and Hopx-CreER;Imp1fl/fl;Rosa-tdTomato mice. C)Quantification of RFP+ cells in Hopx-CreER;Rosa-tdTomato and Hopx-CreER;Imp1fl/fl;Rosa-tdTomato mice. N=4 mice/group, P-value generated by two-tailed student’s t-test. D)Plots demonstrating UEA-1+ cells in Imp1^WT^ and Imp1^ΔIEC^ mice. E)Quantification of UEA-1+ cell numbers in Imp1^WT^ and Imp1^ΔIEC^. N=4 mice/group, P-value generated by two-tailed student’s t-test. F)Plots demonstrating CD24-high cell numbers in Imp1^WT^ and Imp1^ΔIEC^ mice. G)Quantification of CD24-high cell numbers in Imp1^WT^ and Imp1^ΔIEC^ mice. N=4 mice/group, P-value generated by two-tailed student’s t-test. H) Western blot from crypts isolated from indicated mice.

## Acknowledgments

The authors thank Dr. Joseph Baur for helpful discussions and feedback. The authors thank Drs. Sarah Andres and Todd Strochlic for providing plasmid backbones used in the present studies.

## Funding

NIH R01DK124369 (KEH); NIH R21ES031533 (KEH); Institutional Development Funds from Children’s Hospital of Philadelphia Research Institute (KEH); Penn Center for Molecular Studies in Digestive and Liver Diseases NIH P30DK050306 (KEH); NIH R01DK12115 (KAW); NIH DP5OD028144 (SMS); NIH F31-DK124956 (LRP).

## Author Contributions (CRediT Statement)

Study concept and design: Parham, Williams, Hamilton. Acquisition of data: Parham, Williams, Acheampong, Ma, Katada, Simon, Naughton, McMillan, Shaffer. Analysis and interpretation of data: Parham, Williams, Acheampong, Ma, Simon, Danan, Karakasheva, Whelan, Shaffer, Hamilton. Drafting of the manuscript: Parham, Williams, Hamilton. Analysis of the manuscript for intellectual and conceptual content: Parham, Williams, Whelan, Brady, Hamilton. Study Supervision: Hamilton.

## Methods

### Animals

All mice used for these studies were between 8-12 weeks of age, fed ad libitum, and housed under standard ULAR conditions. The following mice were obtained from Jax Laboratories: *Hopx-CreER* (#017606); *Lgr5-GFP* (#008875); *R26-tdTomato* (#007909). *VillinCre;Imp1*^*fl/fl*^ mice were generated previously and maintained on a C57Bl/6 background. Control mice had floxed, intact alleles (*Imp1*^*WT*^). *Atg7-floxed* mice were kindly provided by RIKEN BRC through National Bio-Resource Project of MEXT, Japan [11]. To activate CreERT2, mice received 1mg tamoxifen doses dissolved in corn oil via intraperitoneal injection. For irradiation studies, mice were subjected to 12Gy using the X-RAD 320 Cabinet X-ray System.

### Isolation of small intestinal crypts and FACS analysis

Following mouse euthanasia, the proximal 15cm of the small intestine was isolated and washed in cold PBS. The tissue was splayed open and transferred to a tube containing 10ml of 1X HBSS with 1mM NAC. Following collection, the tissue was vortexed for 15 seconds followed by a 15 second rest on ice; this was performed repeatedly during a 2-minute period. The tissue was then transferred to a tube containing 10ml of 1X HBSS with 1mM NAC and 10mM EDTA and was placed on a rotator in 4°C for 45 minutes. After the incubation period, the tissue was vortexed for 30 seconds followed by a 30 second rest on ice; this was performed repeatedly during a 3-minute period. After vortexing, the tissue digestion was filtered through a 70uM filter and the flow-through was centrifuged at 300g for 3 minutes. To generate a single cell suspension, the cell pellet was resuspended in buffer containing DNAse (35ug/mL) and Liberase (20ug/mL) and was incubated at 37°C for 20 minutes. Following digestion, cells were washed in PBS and were resuspended in media containing the desired antibodies and incubated at 37°C for 30 minutes. The following antibodies and dilutions were used: EpCam-PE (1:500, Thermo Fisher Scientific Cat# 12-5791-82); CD24-APC (1:500, BioLegend Cat# 101813); UEA-1 (1:500, Vector Laboratories Cat# FL-1061). Subsequently cells were washed in PBS and resuspended in FACS buffer (PBS with 4% FBS) prior to FACS analysis. The viability dyes DAPI and DRAQ7 were used to exclude dead cells. Cells were analyzed and sorted on the FACSJazz and FACSAria Fusion sorters. Data analysis was performed using FlowJo software.

### Small intestinal enteroid culture and analysis

Intestinal epithelial cells isolated by FACS were plated in Matrigel droplets at a density of 12,000 cells per 20uL droplet. Matrigel droplets were overlaid with the following medium: advanced DMEM/F12 media containing 1X Glutamax, 10mM Hepes buffer, 1X Antibiotic-Antimycotic, titrated R-spo1 and Noggin-containing conditioned media, 1x N-2 supplement, 1x B-27 supplement, 5uM CHIR99021, 1mM NAC, 50ng/mL mEGF, 5% Noggin/R-spondin conditioned medium, and Y-27632 (10uM). Media was replaced the day after initial plating (Day 1), and every other day afterwards. All organoid experiments were imaged on Day 5 using a Keyence BZ-X all-in-one fluorescence microscope to collect Z-stack images of 3×3 focal planes that were subsequently stitched together to form one image that encompassed the entire Matrigel. The number of organoids per droplet were counted and organoid formation efficiency was calculated as the proportion of organoids relative to total cells plated. Individual experiments were normalized to respective wildtype controls and represented in aggregate to account for inter-experimental variability.

### Live imaging of autophagic vesicles in enteroids

To evaluate autophagic vesicle accumulation in enteroids, CytoID Autophagy Detection Kit was used per manufacturer’s protocol, with modifications. Enteroids were passaged 1 day prior to assays to obtain equivalent plating densities. Three hours prior to imaging, 2 μg/ml CytoID and 2 μg/ml Hoechst33342 (ENZO Life Science) were added in the medium and incubated at 37°C. In chloroquine-treated enteroids, 120 μM of chloroquine (ENZO Life Science) was also added in the medium. After 3 hours incubation, enteroids were washed with PBS and fresh medium was added. Enteroids were immediately analyzed using confocal microscopy ECLIPSE Ti (Nikon).

Six to ten enteroids per mouse were analyzed across three mice per genotype. Additional enteroid photos were obtained using a Leica TCS SP8 Confocal microscope. Genotype-blinded enteroid imaging of CytoID-positive puncta was performed by RM.

### Immunofluorescence staining

Immunofluorescence staining was performed using heat antigen-retrieval in citric acid buffer (pH 6.0) and staining with E-Cadherin (BD Biosciences 610182, 1:1000 dilution). 5-ethynyl-2′-deoxyuridine (EdU) staining was performed using Click-iT® EdU Alexa Fluor® 488 Imaging Kit (C10337) as per manufacturer’s protocol. All sections were stained with DAPI (Invitrogen) and anti-mouse secondary antibodies. No-primary negative controls were also used. Scoring of EdU-positive foci, where a single focus is defined by a cluster of ≥ 5 EdU+ cells from a single clone (colony or hyperproliferative crypt), where quantified across at least 20 high-powered fields per animal for a total of 4 mice per genotype. Blinded scoring was performed by (PC and KEH).

### Generation of HEK293-IMP1-GFP mutant cells

IMP1 deletion was performed using the IGF2BP1 CRISPR/Cas9 knockout plasmid (Santa Cruz Cat# sc-401703) and IGF2BP1 HDR Plasmid (Santa Cruz Cat# sc-401703-HDR) protocol as described by the manufacturer. Cells were selected using 2 ug/mL of puromycin for 2 passages and knockout was confirmed using western blotting. *IMP1-KO* HEK293 cells were used to seed 6-well plates to 60% confluency. Cells were then transfected with pcDNA-GFP, pcDNA-IMP1-GFP, pcDNA-IMP1-S181A-GFP, pcDNA-IMP1-Y396F-GFP, or pcDNA-IMP1-S181A/Y396F-GFP plasmids using lipofectamine 3000 and 2.5 ug of DNA/well for 24 hours in DMEM+10% FBS. Following 24 hours transfection, media was replaced with standard media (DMEM, 10% FBS, Penn/Strep) for 24 hours. Selection with 500 ug/mL G418 occurred for 4-5 days to enrich for a stably expressing population. After selection, cells were sorted for highest 5% GFP expression using Zombie Aqua (BioLegend 423101) and the MoFlo Astrios (Beckman Coulter).

### Western blot

Cells were lysed with lysis buffer (Cell Lysis reagent (Cell Signaling #9803S), Halt Protease Inhibitor Cocktail (Life technologies #78430), 1 mM Sodium Orthovanadate, 10 uM Sodium Fluoride) on ice and spun at 9391 rcf at 4°C for 10 minutes. Running buffer (final concentration 62.5 mM Tris-HCl pH 6.8, 2.5% SDS, 0.002% Bromophenol Blue, 5% B-mercaptoethanol, and 10% glycerol) was added to supernatant and boiled for 6 minutes and loaded onto an SDS-PAGE gel with a percentage of 10%, 12.5%, or 15% acrylamide. Samples were run and transferred onto PVDF membrane using the Bio-rad Trans-blot Turbo system and then blocked in either 5% Milk or BSA in TBS-T (20.7 mM Tris Base, 150.7 NaCl, 0.1% Tween-20, pH 7.6) for 1 hour at room temperature. Primary antibodies were diluted in 5% Milk or BSA in TBS-T and incubated with gentle rocking at 4°C for 24 hours. Blots were washed with TBS-T and then incubated in appropriate secondary antibody for 1 hour at room temperature with rocking. Following washing, blots were incubated with luminol reagent (Santa Cruz Biotechnology sc-2048) per manufacturer protocol and exposed to autoradiography film. All antibodies are listed in Table 1.

**Table 1:**
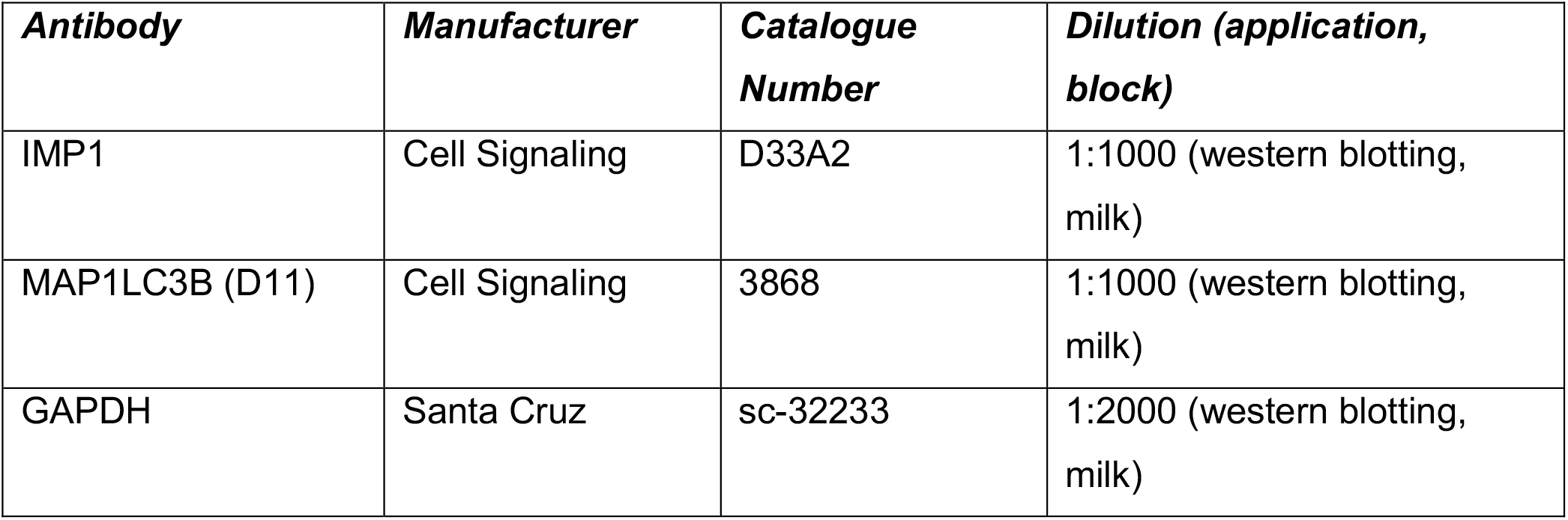
Antibodies

### Single molecule fluorescent in situ hybridization (smFISH)

Single molecule FISH studies were done as described previously [12]. Briefly, cells were grown on coverslips in 6-well plates until confluent. Cells were then fixed with 4% paraformaldehyde for 10 minutes followed by washing and permeabilization in 70% ethanol overnight at 4C. Cells washed in FISH wash buffer (2x SSC, 10% Formamide) then incubated overnight with probes in hybridization buffer (10% dextran sulfate, 10% formamide, 2x SSC) at 37C at dilution of 1:100. After washing, cells stained with DAPI followed by wash in 2x SSC buffer before being mounted using VECTASHIELD Vibrance Antifade Mounting Medium (Vector Laboratories H-1700-2).

Cells imaged using inverted Nikon Ti2-E microscope equipped with a SOLA SE U-nIR light engine (Lumencor), ORCA-Flash 4.0 V3 sCMOS camera (Hamamatsu), with ×60 Plan-Apo λ (MRD01605) or ×20 Plan-Apo λ (Nikon MRD00205). Specific probes are listed in Table 2.

**Table 2:**
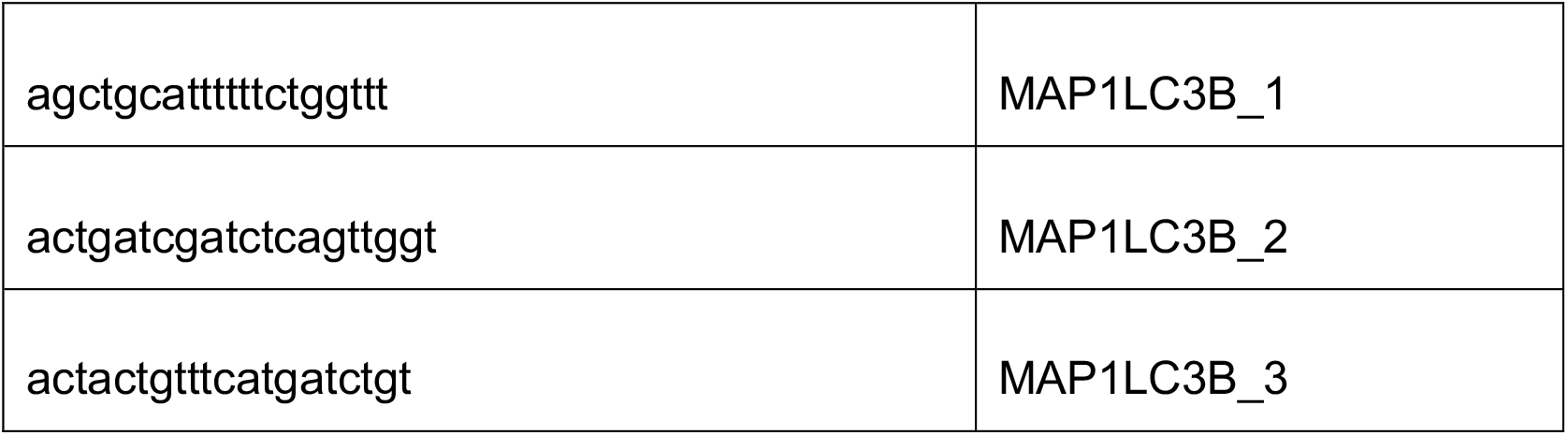
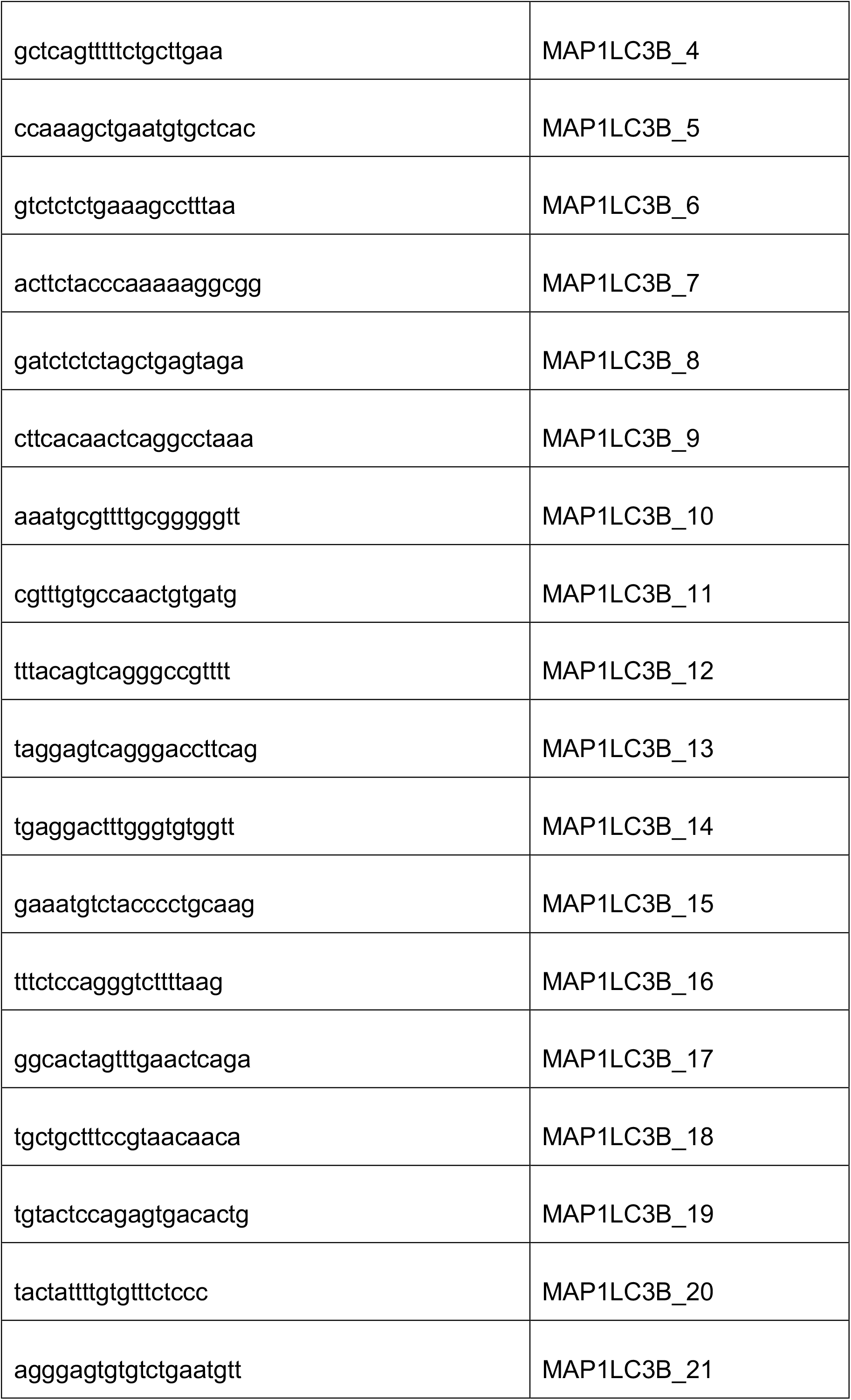
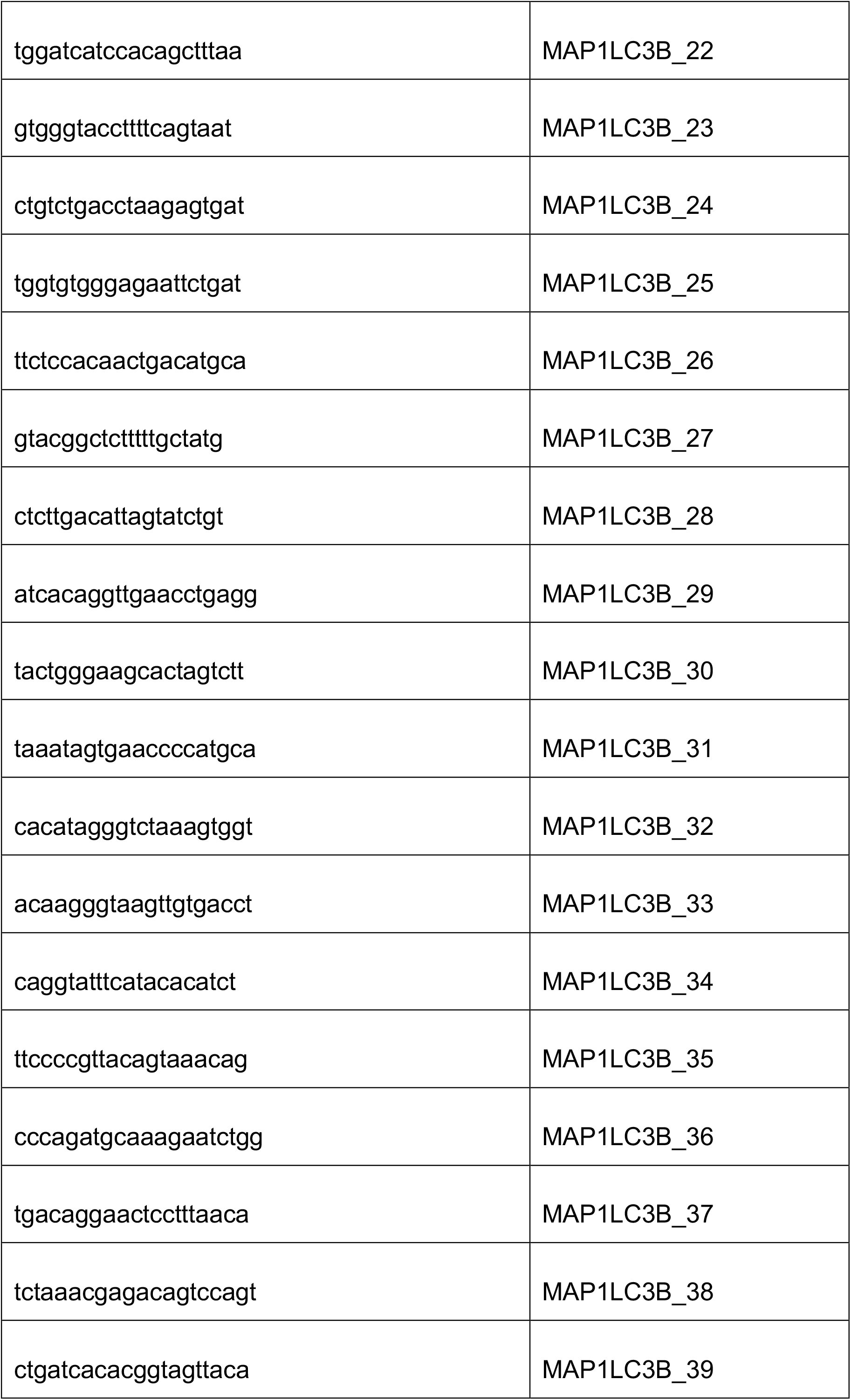
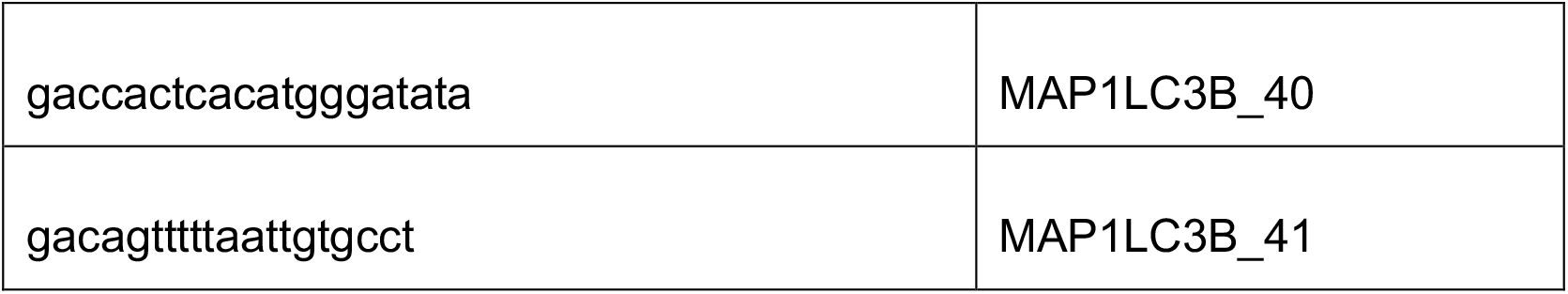
smFISH Probes

Images analyses were performed using ImageJ and CellProfiler [13, 14].

### Statistics

Data were analysed using unpaired, two-tailed student’s t-tests or one-way ANOVA with post-hoc tests (Tukey’s) and p-values indicated in individual figures. Analyses were performed on data from a minimum of three experiments unless otherwise noted, in which case multiple fields are analysed. All values are presented as mean ± standard error of mean (SEM). Specific experimental replicates are described in each figure legend.

## Notes

### Competing Interest Statement

The authors have declared no competing interest.

### Summary of Updates

The manuscript was revised to include additional functional data in mouse models and organoids and text shortened to be submitted as a research letter.

